# Climate warming drives pulsed-resources and disease outbreak risk

**DOI:** 10.1101/2025.01.03.631191

**Authors:** Rémi Fay, Marlène Gamelon, Thibaud Porphyre

## Abstract

Climate influences the risk of disease transmission and spread through its direct effects on the survival and reproduction of hosts and pathogens. However, the indirect influences of climate variation, as those mediated by food resources on host demography, are often neglected. Pulsed-resources produced by oak trees in temperate forests constitute important resources for seed consumers and strongly depend on temperatures. Using an individual-based model, we provide a theoretical exploration of the influence of climate warming on the dynamic of the African swine fever (ASF) in the seed consumer wild boar (*Sus scrofa*), considering both direct and indirect temperature effects. We show that climate warming directly decreases the persistence of the virus in the environment, but also increases the production of acorns with cascading effects on the seed consumer host species. Integrating these climatic effects suggests a decrease of ASF spread under future warmer conditions. Importantly, food-mediated indirect effects of climate may outweigh direct effects, reversing in some situations the predictions of epidemic dynamics under climate change. This shows that anticipating future epidemic risks requires a deep understanding of ecological systems, including all direct and indirect climatic effects.

## 1. Introduction

Climate warming is expected to modify most of the ecosystem processes including disease dynamics. Indeed, because temperature optima generally differ between hosts and pathogens, climate warming is expected to modify host-pathogen interactions (Cohen *et al*. 2020; Thomas & Blanford 2003). Although anticipating the influence of climate warming on hosts and pathogens dynamics is critical to deliver effective prevention measures to mitigate future epidemic risks, this is challenging as temperature impact their dynamics both directly and indirectly.

Temperature directly influences the demographic rates of hosts and pathogens, i.e. their survival, reproduction, infectiousness (Johnson *et al*. 2007; Marcus *et al*. 2023; Paniw *et al*. 2022). As a result, contact rates among hosts and pathogen transmission may change, thereby changing the risk of disease spread and epidemic outbreak. Many studies have assessed the direct effects of temperature on hosts and pathogens, determining for instance thermal performance curves in experimental studies (Ferguson & Adamo 2023; Tadiri & Ebert 2023). Hosts-pathogens dynamics can also be influenced by indirect temperature effects, particularly those that are mediated by food resources. Changes in food availability can impact host demographic rates (Becker *et al*. 2015; Johnson *et al*. 2007), movements and aggregation (Merkle *et al*. 2018), and immune defenses by strengthening or weakening allocation trade-off among competing physiological functions (Martin *et al*. 2008). Temperature effects on food resources could be a major route, especially for homeotherm hosts such as mammals, which are likely less vulnerable to direct temperature effects than ectotherms. Although they may be important drivers in disease spread, indirect temperature effects are often overlooked when assessing future risk of disease epidemics in natural populations.

Some pulsed resources produced by plants, such as acorns produced by oak trees, are driven by temperatures and are thus expected to change under climate warming (Bogdziewicz *et al*. 2021; Schermer *et al*. 2019). These mast events which lead to strong annual variation of seeds or fruits abundance over large areas, have a strong impact on animal consumers being potential reservoirs for disease (e.g. rodents, wild boar) (Ostfeld & Keesing 2000). Therefore, a system involving a pathogen and a pulse resource consumer as hosts constitutes an excellent case study to investigate direct and indirect effects of temperatures on epidemic dynamics in natural populations.

Here, we assess how climate warming directly and indirectly influences host-pathogen dynamics using the study case of the wild boar *Sus scrofa* and the African swine fever (ASF), a highly contagious viral disease that is currently spreading worldwide. The virus can spread through contacts among living wild boars, and through contacts with infected carcasses (Sauter-Louis *et al*. 2021). In this system, increasing temperatures have direct negative effects on virus persistence in the environment (Mazur-Panasiuk *et al*. 2019), and indirect effects on wild boar, through positive influence on oak tree masting (Schermer *et al*. 2019), a key food resource for this host species. High acorn production decreases foraging movements (Morelle *et al*. 2015) and thus opportunities to contact carcasses (Ballari & Barrios-García 2014; Cukor *et al*. 2020; Probst *et al*. 2017), advances wild boar birth peak (Cachelou *et al*. 2022), and increases female breeding proportion (Gamelon *et al*. 2021b) (Figure 1).

**Figure 1:**
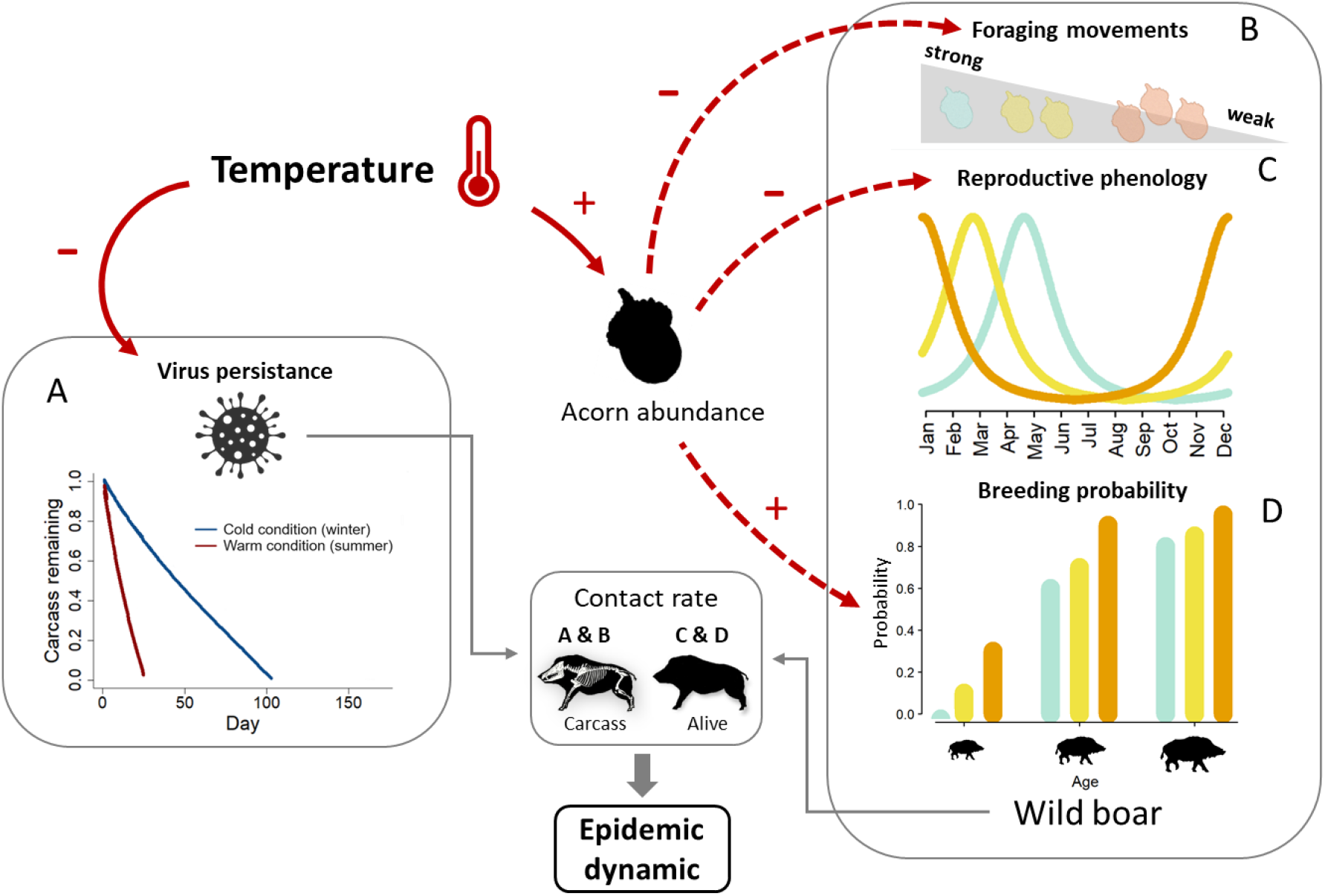
Overview of the influence of the temperature on the wild boar and ASF dynamics. Solid red arrows figure direct temperature effects and dashed red arrow figure indirect temperature effects mediated through acorn production, with the sign indicating the positive or negative influence of temperature. (A) Carcass degradation as a function of temperature driving virus persistence in the environment (illustration for a 40 kg carcass with average winter and summer temperature). (B) Foraging movements according to acorn abundance (blue=low, golden=medium, orange=high). (C) Reproductive phenology (birth peak) according to acorn abundance. (D) Variation in age-specific breeding probability according to acorn abundance. All these processes affect contact rates, both among alive individuals and with carcasses, impacting epidemic dynamics.

Capitalizing on a long-term individual monitoring of a wild boar population in France, along with temperature and acorn production records on the same area, we developed a stochastic individual-based model (IBM) to describe the daily interplay between wild boar population dynamics and ASF transmission dynamics. Our study population is free from this virus and thus epidemic dynamics were investigated using simulations, with virus introduction at variable months to account for seasonal variation (Fay *et al*. 2024). We evaluated both direct and indirect effects of temperature on the wild boar-ASF epidemic dynamics, assessing the contribution of each pathway. Then, we projected host-pathogen dynamics under the RCP 8.5 climate warming scenario to anticipate potential epidemic changes in the coming decades.

## 2. Material and Methods

### 2.1 Study system

The wild boar is a large omnivorous, polygynous and sexually dimorphic ungulate widespread in Eurasia (Massei *et al*. 2015). This study focuses on a hunted population monitored since 1982 in the forest of Châteauvillain-Arc-en-Barrois (48°02N, 4°56E), north-eastern France. This population occupies forests dominated by oaks (*Quercus petraea*) and lives under climatic condition intermediate between continental and oceanic. The demography of this population is highly seasonal with a birth peak in spring and a high mortality rate during the hunting season (Oct-Feb) (Gamelon *et al*. 2021a). A capture-mark-recapture program is running allowing data on wild boar demographic rates to be collected since the 1980’s (Table S1). Annual acorn production was measured indirectly by analyzing stomach contents of shot individuals. Three categories of acorn production (high, medium and low) were defined depending on the quantity of seeds found in the stomachs (for further details see Servanty *et al*. (2009)). Finally, average daily air temperature was measured from 2003 to 2022 on six Météo France weather stations surrounding our study area (<50km).

The ASF virus is a DNA-virus of the Asfarviridae family which infects suids species only. In 2007, the disease was introduced in Europe and rapidly spread both Westward and Eastward, and is now covering most of Europe and Asia (Sauter-Louis *et al*. 2021). Susceptible individuals can be infected through close contacts with infectious individuals (nose to nose contacts, saliva, urine), or with infected carcasses that remain in the environment. With a case-fatality rate approaching 100% and no vaccine available, ASF represents a major socio-economic burden and a threat to food security and biodiversity (Sauter-Louis *et al*. 2021). Our study population is not yet infected by the ASF.

### 2.2 Direct and indirect temperature effects on the wild boar – ASF system

Temperature has both direct and indirect effects on this system. Temperature directly modulates ASF persistence in the environment by affecting carcass persistence. ASF survives in a carcass as long as it persists, that is for months in cold winter conditions and a few weeks under warm summer conditions (Matuszewski *et al*. 2014) (Figure 1A). In contrast, while there is no evidence for a direct effect of temperature on wild boar’s demographic rates in our study population, indirect effects exist through oak mast. Oak mast is highly dependent on thermal conditions during the pollen emission period, and warmer spring conditions increase the frequency of mast seeding events (i.e. massive fruit production spatially synchronized among trees) (Schermer *et al*. 2019). Acorn constitutes the preferred food resources for wild boars, and their abundance influence they foraging movements. Especially, when acorn abundance is low, wild boars increase their foraging movement (Morelle *et al*. 2015) and thus their probability to meet and contact a carcass (Figure 1B). High acorn abundance also increases the breeding probability of females and leads to the advance of their reproductive phenology (Figure 1C,D), but does not affect survival probabilities (Cachelou *et al*. 2022; Gamelon *et al*. 2021b; Touzot *et al*. 2020). While increasing food abundance may improve body condition, which may increase immune defense and survival (Becker *et al*. 2015), ASF is a highly virulent pathogen and we assumed that immune system efficiency is unlikely to be changed, as suggested by the high mortality rate of captive infected individuals which enjoyed *ad libitum* food (Gallardo *et al*. 2017). Because oaks are abundant and homogeneously spread over our study area, we assumed that masting events do not modify host aggregation.

### 2.3 Modelling framework

We expand a stochastic IBM previously developed to describe the daily interplay between this wild boar population and ASF transmission dynamics (Fay et al. 2024). A detailed presentation of the model is provided in Supplementary Information. Briefly, the model is composed of two major components, representing (i) the demographic dynamics of the host population, and (ii) the transmission dynamics of the virus within the population. Hosts were modeled individually and characterized by sex, age and health status. Each day, individuals may successively survive or die, breed or not, and change health status (Table S1 for parameter values). Wild boar reproduction rate, phenology and contact rate with carcasses were dependent on acorn abundance (Table S1). The infection dynamics in the population was modelled considering an SEIR epidemiological process, in which individuals are either susceptible (S), exposed but not infectious (E), infectious (I) or recovered (R). Individuals that died overtime, either from natural causes or from the disease, remain as carcasses in the environment. As such, susceptible individuals can be infected through contacts with alive infectious individuals (density dependent) or with infected carcasses (frequency dependent). All animals killed during the hunting season were removed from the population and thus do not participate in the disease transmission dynamics. Finally, this IBM is not spatially explicit and we assume that we modelled a localized and closed wild boar population, corresponding to situations where a population has been isolated following disease detection, as done in Europe (Sauter-Louis *et al*. 2021). Because oaks are abundant and homogeneously spread in our study area, assuming no spatial heterogeneity is valid for this food resource.

Each simulation run was started assuming a population of 100 individuals, mimicking a small and localized wild boar population. Initial age and sex distributions were attributed according to the stable age and sex distribution. All individuals were immunologically naïve (susceptible). Host population was projected one year prior to the introduction of the virus to insure independence from the initial conditions. The virus was introduced the second year in the population by adding one infected sub-adult male (age = 1.5 years), illustrating the immigration of an infected individual within the population. Because the timing of virus introduction has strong influence on the subsequent epidemic dynamics (Fay *et al*. 2024), each model was run 12 times introducing the virus on the first day of each month. Following virus introduction, the population was projected for two additional years, leading to a total projection length of four years.

### 2.4 Assessing temperature effect on the ASF-wild boar system

#### 2.4.1 Epidemic dynamics under current climatic conditions

We characterized the epidemic dynamics under current conditions based on temperature and acorn production time-series observed from 2003 to 2022. We first repeatedly run the model for all potential sequences of four consecutive years for temperature and corresponding acorn abundance. These models allow us to depict the influence of inter-annual temperature variation, integrating the direct and indirect temperature effects on epidemic dynamics. To assess the epidemic dynamic under current climatic condition, we run the model using the average daily temperature profile from 2003 to 2022, and sampled random sequences of four consecutive years of acorn abundance.

Then, we run additional models to disentangled the direct and indirect effects of temperature. To evaluate the direct effect of temperature, we run a set of models, each model considering a specific series of annual temperature profile measured over the study period. For each model, acorn abundance sequences were randomly sampled from the observed acorn production series to control for indirect temperature effects. To assess the indirect effects of temperature on epidemic dynamics, we run three models corresponding to the three levels of acorn abundance, while using the average daily temperature variation to control for the direct temperature effect. Because indirect effects operate through three pathways (i.e. influence carcass attractivity, reproductive phenology and female breeding probability, Figure 1B, C, D), we also run models including only one pathway at a time to assess their relative effects.

#### 2.4.2 Epidemic dynamics under future climatic conditions

Expected future temperatures in the study area were projected until 2100 using five alternative climatic models from the EURO-CORDEX initiative (Kotlarski *et al*. 2014), assuming the RCP 8.5 scenario of future greenhouse gas emission. This “business as usual” scenario is the most pessimistic leading to end-of-century warming outcomes from 3.3 °C to 5.4 °C (5th to 95th percentile) with a median of 4.5 °C. While this scenario may overestimate gas emission by 2100, its predictions match current emissions and are judged realistic for the midcentury (Schwalm *et al*. 2020). Acorn projections corresponding to these temperature projections were obtained using a mechanistic resource budget model linking temperatures to oak production (Schermer *et al*. 2019; Touzot *et al*. 2020). Expected acorn production, first obtained on a continuous scale, were discretized according to the three levels of acorn abundance (i.e., low, medium, high). Finally, we projected the wild boar–ASF system under expected future climate conditions considering two-time windows: 2050-2069 and 2080-2099. This allowed us to project this system under a lower and upper range of warming. We run the model using the simulated sequences of temperatures and acorn abundance and compared the average epidemic dynamics to estimates obtained from the current climatic conditions. As mentioned earlier, we run additional models to disentangled the contribution of direct and indirect effects of temperature in the future climatic conditions.

### 2.5 Model outputs

The model was built and run using R statistical software version 4.2.2 (R Core Team 2022). Each scenario was run multiple times to observe 1,000 successful virus invasions, allowing us to describe the epidemic dynamics accounting for stochasticity in model output. To describe the epidemic dynamics, we computed the basic reproduction number (*R*_*0*_), that is the number of secondary infections generated by a single infected individual. Values of *R*_*0*_ above one indicate that pathogen will grow and generate outbreaks, while values of below suggest the opposite. We also estimated the invasion success, here defined as the probability that at least five individuals were infected. Then, given invasion success, we computed the maximum daily incidence rate (i.e., maximum number of new cases observed in a single day) and the epidemic duration.

## 3. Results

Interannual variation of temperature generates strong variation in epidemic dynamics (Figure 2). Relative differences from the average pattern were positive or negative according to the year and reached 14% for the basic reproduction number (i.e., the mean number of secondary cases generated by a given infectious case in an entirely susceptible population, *R*_0_) and the invasion success (i.e. the probability single incursion generates outbreaks of ≥5 cases), 17% for the daily maximum incidence rate and 15% for epidemic duration (Table S2). While both direct (via virus persistence) and indirect (via food resource) temperature effects generate this interannual variability, direct temperature effects contributed less to the observed variation in epidemic dynamics than indirect effects: 6% vs. 14% for R_0_, 4% vs. 15% for invasion risk and 3% vs. 15% for the maximal incidence rate (Figure 3, Table S1). The only exception was for the epidemic duration that tended to be more affected by the direct effects than by indirect effects (16% vs. 11%, Table S2). Influence of temperature on epidemic dynamics varied seasonally according to the timing of virus introduction, especially for the maximum incidence rate that was more affected by temperature in summer and autumn (June-November, 20%), than in winter and spring (December-May, 12%) (Figure 2, Table S2). Finally, disease seasonality showed moderate variation according to the thermal condition (Figure 2). Seasonal pattern of the epidemic dynamics was poorly influenced by direct temperature effects, but more strongly by indirect temperature effects. Under high acorn abundance, virus invasion success was higher and subsequent outbreaks larger in case of late winter virus introduction (February-March) compared to late spring virus introduction (May-June), but the reverse pattern was observed under low acorn abundance (Figure 3).

**Figure 2:**
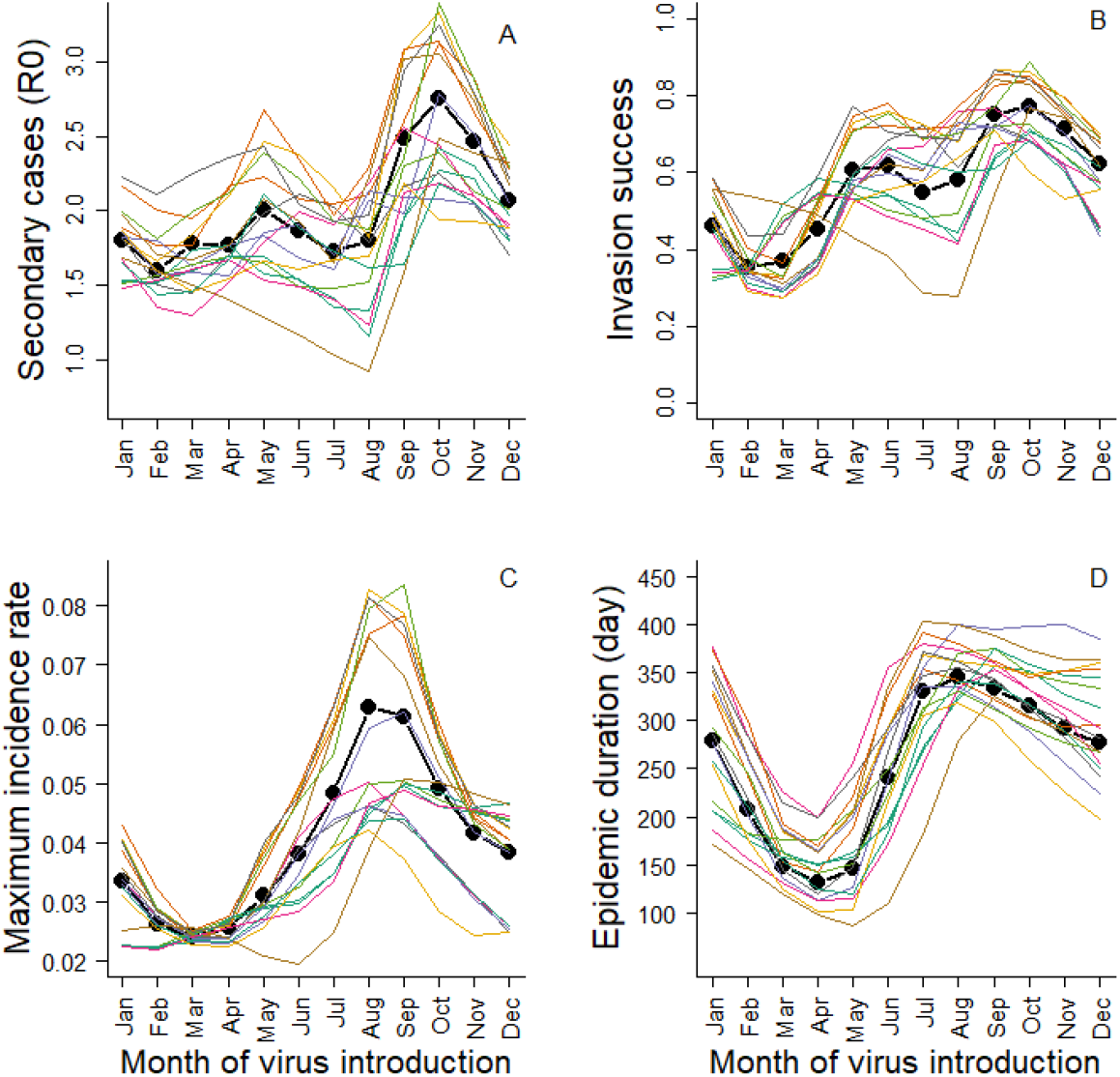
Epidemic dynamics variation generated by interannual variation in temperature between 2003 and 2022 (colored lines) (both direct and indirect effects of temperature are considered). The bold black line shows the epidemic dynamic under the average environmental conditions. Results include (A) the basic reproduction number (*R*_*0*_), (B) the invasion success probability defined as the probability that there were at least five secondary cases, (C) the maximum daily incidence rate and (D) the epidemic duration conditional on successful virus invasion. Results are presented as a function of time of virus introduction in the population from 1^st^ January to 1^st^ December.

**Figure 3:**
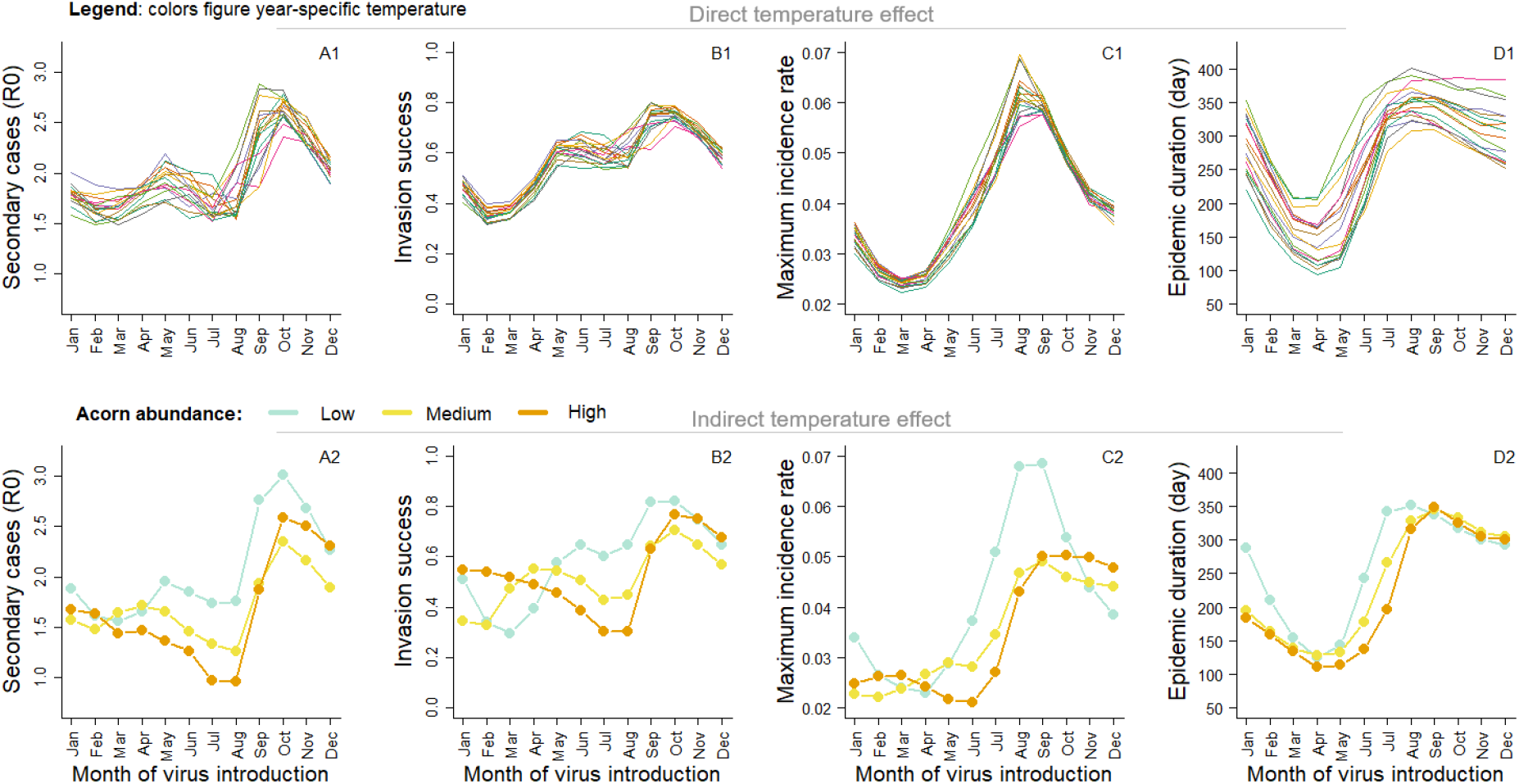
Direct (i.e. through virus persistence in the environment) and indirect (i.e. through acorn abundance) effect of temperature on epidemic dynamic. Results include (A) the basic reproduction number (*R*_*0*_), (B) the invasion success probability defined as the probability that there were at least five secondary cases, (C) the maximum daily incidence rate and (D) the epidemic duration conditional on successful virus invasion. Results are presented as a function of time of virus introduction in the population from 1^st^ January to 1^st^ December.

Projecting this host-pathogen system in warmer conditions at the end of the century under the RCP 8.5 scenario, we found that epidemic dynamics were likely to be weaker, with lower *R*_0_ (-13%), lower invasion success (-8%), lower maximum daily incidence rate (-13%) and shorter duration (-21%) relatively to current temperature conditions (Figure 4, Table 1). Both direct and indirect effects led to lower epidemic dynamics, but direct temperature effects contributed more to the future change in epidemic dynamics than indirect effects (average for 2080-2099: -13% vs. -3%, Figure S3 & S4). Importantly, changes in future epidemic dynamics were season-specific (Figure 4). Negative changes were stronger when the virus was introduced in summer and autumn (range: -18% to -21%) than in winter and spring (range: -5% to +5%), though epidemic duration remains relatively stable across seasons (Table 1). Furthermore, some changes were reversed according to the seasonality. For instance, invasion success was predicted to decrease over summer and autumn condition (-18%), but to increase in autumn and winter (+5%) (Table 1).

**Table 1:** Projected epidemic change under future temperature conditions. Estimates were obtained by comparing the epidemic dynamics under average current temperature conditions to models figuring future climate conditions, including either the influence of temperature through both direct and indirect effects (Direct & Indirect), direct effects only (Direct) or indirect effects only (Indirect). Results include the average change in the focus epidemic metric over the entire year (Full year), and then distinguishing two periods within a year: winter and spring (December-May), and summer and autumn (June-November).

|  |  | % change compared to current temperature conditions |  |  |  |  |  |
| --- | --- | --- | --- | --- | --- | --- | --- |
|  |  | Full year |  | Dec-May |  | Jun-Nov |  |
|  | Temperature pathway | 2050-2079 | 2080-2099 | 2050-2079 | 2080-2099 | 2050-2079 | 2080-2099 |
| **$R_0$** | Direct & Indirect | -5 | -13 | +3 | -5 | -11 | -21 |
|  | Direct | -4 | -13 | -1 | -8 | -7 | -17 |
|  | Indirect | -1 | -3 | +5 | +5 | -5 | -9 |
| **Invasion success** | Direct & Indirect | -3 | -8 | +4 | +5 | -9 | -18 |
|  | Direct | -3 | -9 | -2 | -9 | -4 | -9 |
|  | Indirect | +0.4 | -0.2 | +8 | +11 | -5 | -8 |
| **Maximum incidence** | Direct & Indirect | -9 | -13 | -1 | -4 | -13 | -19 |
|  | Direct | -5 | -11 | -4 | -7 | -6 | -13 |
|  | Indirect | -5 | -6 | +1 | +2 | -8 | -10 |
| **Epidemic duration** | Direct & Indirect | -13 | -21 | -15 | -23 | -11 | -20 |
|  | Direct | -11 | -20 | -13 | -22 | -10 | -19 |
|  | Indirect | -2 | -2 | -2 | -4 | -2 | -1 |

**Figure 4:**
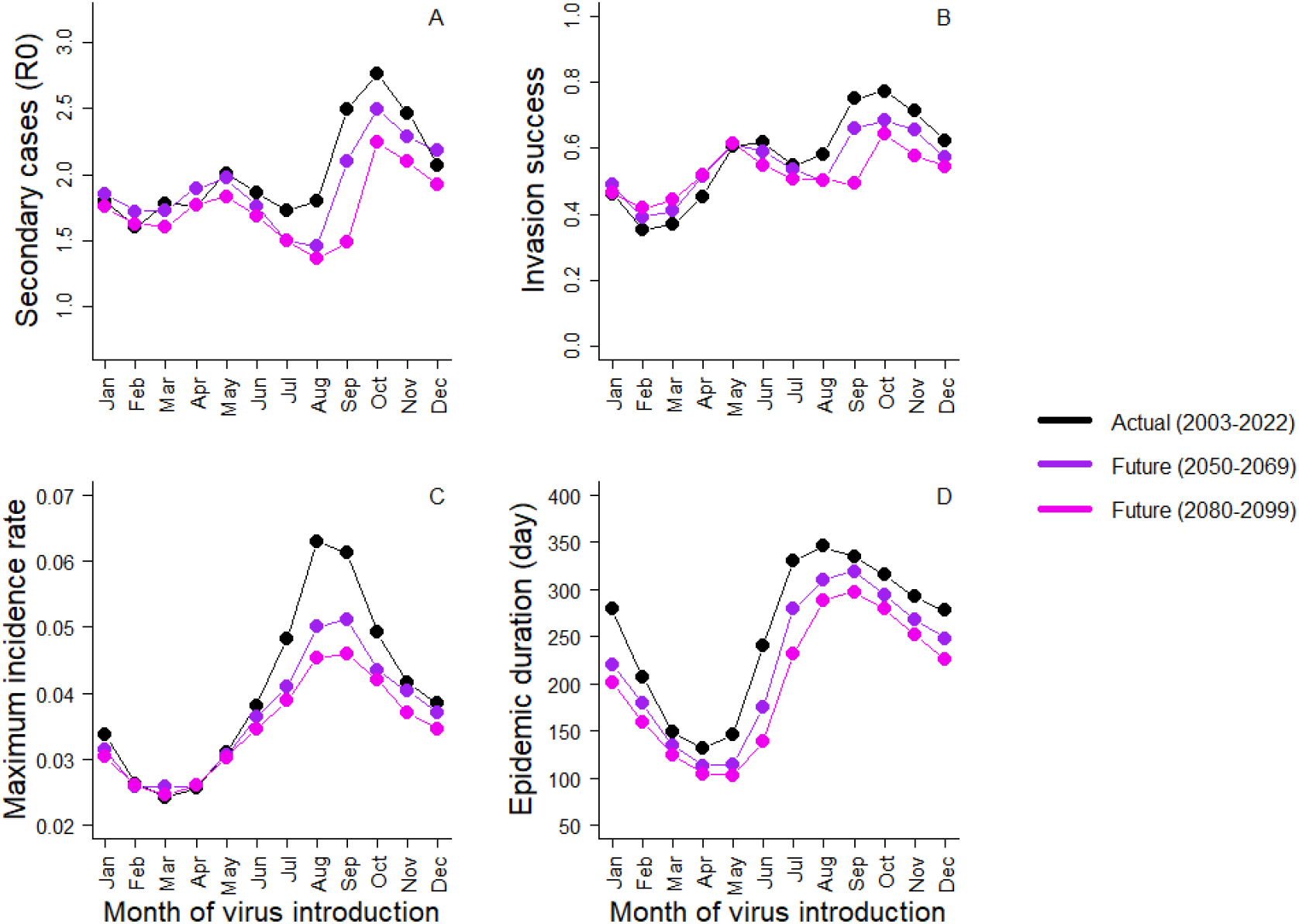
Total influence (i.e. including both direct and indirect effects) of current (black) and future (purple, pink) temperature on epidemic dynamics. Results include (A) the basic reproduction number (R0), (B) the invasion success probability defined as the probability that there were at least five secondary cases, (C) the maximum daily incidence rate and (D) the epidemic duration conditional on successful virus invasion. Results are presented as a function of time of virus introduction in the population from the 1^st^ January to 1^st^ December.

## 4. Discussion

Acorn abundance determines wild boar reproductive phenology (Cachelou *et al*. 2022) with consequences on host-pathogen phenological match and disease seasonality (Figure S1). On the one hand, there are higher contact rates and thus pathogen spread among individuals during the birth peak, when both population size and the proportion of young individuals peak. On the other hand, pathogen persistence is higher in carcasses at low temperatures. As warmer springs favor high acorn production, push forward the birth peak of wild boar and decrease virus persistence in the environment, the risk of virus invasion decreases and outbreaks are shorter and less intense (Figures 3, 4). Phenological mismatch between consumers and resources has been frequently reported as a consequence of climate change (Roslin *et al*. 2021), with potential negative impacts for consumer demography (Visser & Gienapp 2019). Here, we show that similar mismatch may occur in a host-pathogen system. As a change in host phenology in response to climate change is likely to be frequent along with changes in pathogen seasonality (Gethings *et al*. 2015; McDevitt-Galles *et al*. 2020; Paull & Johnson 2014; Tadiri & Ebert 2023), changes in the timing of host–pathogen interactions could be widespread in the wild. Thus, temperature-induced changes in host and pathogen phenology, rather than changes in the total abundance of host or pathogen alone, can be crucial for determining how temperature change will influence epidemic dynamics (Gethings *et al*. 2015; Paull & Johnson 2014).

In the wild boar–ASF system, indirect temperature effects mediated through oak masting is a major pathway by with temperature influences epidemic dynamics. In other host-disease contexts, fluctuations in acorn abundance positively influence rodent population demography and Lyme disease prevalence (Ostfeld *et al*. 2006). Similarly, changes in grass phenology has been shown affecting elk movements and aggregation, with consequences on brucellosis transmission risk (Merkle *et al*. 2018). Indirect temperature effect could be even more complex. In a snail species *Helisoma trivolvis*, warm conditions were found to increase food availability, which, as a consequence, promotes body growth and increases host survival. This improve both host tolerance to infection and parasite (*Ribeiroia ondatrae*) reproduction rate within infected hosts, which lead ultimately to higher parasite prevalence (Paull & Johnson 2014). Changes in food resources under current global warming could thus be an important ecological pathway by which climate influences disease dynamics indirectly in the wild.

Because epidemic responses to climate warming result from the direct and indirect effects of temperature on hosts and pathogens, considering only one component (i.e., host or pathogen, direct or indirect temperature effect) could lead to biased predictions (Paniw *et al*. 2022). In the case of ASF, epidemic risk was predicted lower over winter and spring when considering direct effect only, but the opposite was expected after integrating both direct and indirect effects (Figure 4, Figures S2 & S3). Similar shift in predictions when integrating or not all temperature effects has been reported in other systems. For instance, in a caterpillar, temperature increases feeding rate and such exposure to a food-mediated pathogen (baculovirus), suggesting an increase of disease prevalence under warmer conditions (MacDonald *et al*. 2023). Yet, increasing temperatures also increase the virulence of the pathogen leading to the quicker death of infected hosts which decrease exposure of susceptible individuals. The latter effect counterbalanced the former, making the disease prevalence lower in a warmer environment (MacDonald *et al*. 2023).

Host and pathogen dynamics will inevitably change with climate change, creating potentially new ‘winners’ and ‘losers’ in these interactions (Thomas & Blanford 2003). However, depending on the study system, winner could be either the host (Gethings *et al*. 2015; Paull & Johnson 2014) or the pathogen (Paniw *et al*. 2022; Tadiri & Ebert 2023). The output of the new host-pathogen interaction will depend on the system and ecological context, with potentially spatially heterogenous response for the same host-pathogen system (Gethings *et al*. 2015). Overall, predicting the influence of climate change on disease outbreak requires detailed knowledge on the system, emphasizing the value of a deep understanding of ecological systems as a whole.

## Data accessibility statement

Code from this study has been deposited in Zenodo repository https://doi.org/10.5281/zenodo.10605216

## Acknowledgments

This work was funded by the French National Research Agency and Boehringer Ingelheim Animal Health France through the IDEXLYON project (ANR-16-IDEX-0005) and the Industrial Chair in Veterinary Public Health, as part of the VPH Hub in Lyon. We thank the Office Français de la Biodiversité (OFB) and all the people involved in data collection. We are particularly grateful to Eric Baubet for its valuable comments on this manuscript. This study is part of the long-term Studies in Ecology and Evolution (SEE-Life) program of the CNRS. This work was performed using the computing facilities of the CC LBBE/PRABI.

## Authors’ contributions

All authors conceived the ideas. RF performed the modeling, analyses and interpretation with the help of MG and TP. RF wrote the paper with feedback and editing from all co-authors.

