## Supplementary materials for "Climate warming drives pulsed-resources and disease outbreak risk"

**In this document:**

**Table S1:** Input parameter values used in a stochastic individual based (p. 2)

**Table S2:** Contribution of direct and indirect temperature effects in generating interannual variation in epidemic dynamics (p. 3)

**Figure S1:** Indirect effects of temperature on epidemic dynamics through the three pathways (p. 4)

**Figure S2**: Projection of the epidemic dynamics in future climatic conditions including only direct temperature effects (p. 5)

**Figure S3**: Projection of the epidemic dynamics in future climatic conditions including only indirect temperature effects (p. 6)

**Appendix S1**: Model description following the ODD protocol (p. 7)

**References** (p. 12)

**Table S1**

Input parameter values used in a stochastic individual based model to describe wild boar population dynamics along with the ASF transmission.

| Parameter | Distribution and value | Reference |
|---|---|---|
| **Wild boar demography** | | |
| Daily survival probability: | | |
| Piglet (≤ 1 month) | Bernoulli, p = 0.9968437 | |
| small female (hunting season) | Bernoulli, p = 0.9903375 | |
| medium female (hunting season) | Bernoulli, p = 0.9958889 | |
| large female (hunting season) | Bernoulli, p = 0.9976827 | |
| small female (no hunting) | Bernoulli, p = 0.9999162 | |
| medium female (no hunting) | Bernoulli, p = 0.9993259 | |
| large female (no hunting season) | Bernoulli, p = 0.9993744 | (1, 2) |
| small male (hunting season) | Bernoulli, p = 0.9903375 | |
| medium male (hunting season) | Bernoulli, p = 0.9956167 | |
| large male (hunting season) | Bernoulli, p = 0.9900668 | |
| small male (no hunting) | Bernoulli, p = 0.9998306 | |
| medium male (no hunting) | Bernoulli, p = 0.9989161 | |
| large male (no hunting season) | Bernoulli, p = 0.9995629 | |
| Annual reproduction probability*: | | |
| small female (acorn=low) | Bernoulli, p = 0 | |
| medium female (acorn=low) | Bernoulli, p = 0.60 | |
| large female (acorn=low) | Bernoulli, p = 0.80 | |
| small female (acorn=medium) | Bernoulli, p = 0.10 | |
| medium female (acorn=medium) | Bernoulli, p = 0.70 | (3) |
| large female (acorn=medium) | Bernoulli, p = 0.85 | |
| small female (acorn=high) | Bernoulli, p = 0.30 | |
| medium female (acorn=high) | Bernoulli, p = 0.90 | |
| large female (acorn=high) | Bernoulli, p = 0.95 | |
| Reproductive phenology (see Figure 1) | | |
| Birth peak (acorn=low) | 1st May | |
| Birth peak (acorn=medium) | 1st March | (4) |
| Birth peak (acorn=high) | 1st January | |
| Litter size: | | |
| small female (1-9 piglets) | Multinomial distribution, p = 0.034, 0.132, 0.231, 0.267, 0.199, 0.098, 0.032, 0.007, 0.001 | |
| medium female (1-10 piglets) | Multinomial distribution, p = 0.009,0.056,0.133,0.226,0.256, 0.189,0.091,0.031,0.008,0.001 | (5) |
| large female (2-13 piglets) | Multinomial distribution, p = 0.003,0.015,0.046,0.105,0.180, 0.222,0.199,0.132,0.066,0.024, 0.006,0.001 | |
| Litter sex-ratio | Binomial, mean = 0.5 | (6) |
| **ASF epidemiology** | | |
| Daily number of contacts with: | | |
| relative | fct(A=0, K=10, r=0.03, $v$=0.1, $\gamma_{age}$=3, $\gamma_{sex}$=4, $f_{t,sex_i}$=20) | (5) |
| carcass (acorn=low) | Linear model, $\delta$ = 0.012 | Assumption reflecting the expected |
| carcass (acorn=medium) | Linear model, $\delta$ = 0.006 | decrease of contact rate with carcasses with |
| carcass (acorn=high) | Linear model, $\delta$ = 0.003 | acorn abundance (appendix S1) |
| AFS transmission probability per contact with | | |
| relative | 0.05 | (5) |
| carcass | 0.01 | |
| Incubation period (day) | 4 | |
| Duration of infectious period (day) | Poisson, mean = 5, truncated at 1 | (7) |
| Probability of death following infection | 0.95 | |

*We reported annual reproductive probabilities for convenience but daily breeding probabilities were used in the model (5)

**Table S2:** Importance of direct (via virus persistence) and indirect (via food resource) temperature effects, including the three indirect pathways, in generating interannual variation in epidemic dynamics. Changes in the epidemic metrics were obtained by comparing the epidemic dynamic under average current temperature conditions to six models considering either both direct and indirect temperature effect (*Direct & Indirect*), only direct temperature effect (*Direct*), only indirect temperature effect (*Indirect*), or only one indirect pathway (*Carcass attractivity, Reproductive phenology, Breeding probability*). Results include both average change (+/-) or absolute average change (abs) change in the focus epidemic metric over the entire year (Full year), and then distinguishing two periods within a year: winter and spring (December-May), and summer and autumn (June-November).

| | | % of average variation | | | | | |
|---|---|---|---|---|---|---|---|
| | **Temperature pathway** | **Full year** | | **Dec-May** | | **Jun-Nov** | |
| | | **+/-** | **abs** | **+/-** | **abs** | **+/-** | **abs** |
| ***$R_0$*** | Direct & Indirect | -2 | 14 | -1 | 12 | -3 | 16 |
| | Direct | -3 | 6 | -1 | 5 | -4 | 7 |
| | Indirect | -10 | 14 | -6 | 10 | -13 | 16 |
| | Indirect (Carcass attractivity) | -19 | 21 | -23 | 27 | -14 | 17 |
| | Indirect (Reproductive phenology) | -5 | 18 | +7 | 15 | -18 | 20 |
| | Indirect (Breeding probability) | +11 | 12 | +14 | 12 | +8 | 13 |
| **Invasion success** | Direct & Indirect | 0 | 14 | -1 | 15 | -0.1 | 14 |
| | Direct | -1 | 5 | -1 | 5 | -2 | 5 |
| | Indirect | -5 | 15 | +2 | 16 | -11 | 15 |
| | Indirect (Carcass attractivity) | -17 | 21 | -26 | 33 | -7 | 12 |
| | Indirect (Reproductive phenology) | -2 | 21 | +12 | 24 | -17 | 20 |
| | Indirect (Breeding probability) | +10 | 13 | +15 | 13 | +6 | 13 |
| **Maximum incidence** | Direct & Indirect | -3 | 17 | -1 | 12 | -4 | 20 |
| | Direct | -1 | 4 | 0 | 4 | -1 | 4 |
| | Indirect | -8 | 15 | -3 | 11 | -13 | 18 |
| | Indirect (Carcass attractivity) | -17 | 20 | -12 | 16 | -23 | 23 |
| | Indirect (Reproductive phenology) | -5 | 14 | +2 | 10 | -12 | 17 |
| | Indirect (Breeding probability) | +13 | 14 | +21 | 9 | +5 | 17 |
| **Epidemic duration** | Direct & Indirect | +5 | 15 | +8 | 21 | +3 | 12 |
| | Direct | +6 | 12 | +8 | 18 | -4 | 9 |
| | Indirect | -6 | 11 | -6 | 13 | -6 | 10 |
| | Indirect (Carcass attractivity) | -9 | 10 | -9 | 13 | -9 | 9 |
| | Indirect (Reproductive phenology) | -3 | 9 | -2 | 9 | -4 | 9 |
| | Indirect (Breeding probability) | +6 | 7 | +5 | 10 | +8 | 5 |

**Figure S1**: Indirect effects of temperature on epidemic dynamics distinguishing the influence of acorn abundance (blue=low, golden=medium, orange=high) through three pathways: carcass attractivity, reproductive phenology and breeding probability. We used low acorn abundance as a reference scenario for fixed parameters so that the blue line could be used as a reference curve. Results include the basic reproduction number ($R_0$), the invasion success probability, the maximum daily incidence rate and the epidemic duration conditional on successful virus invasion. Results are presented as a function of time of virus introduction in the population from the 1st January to 1st December.

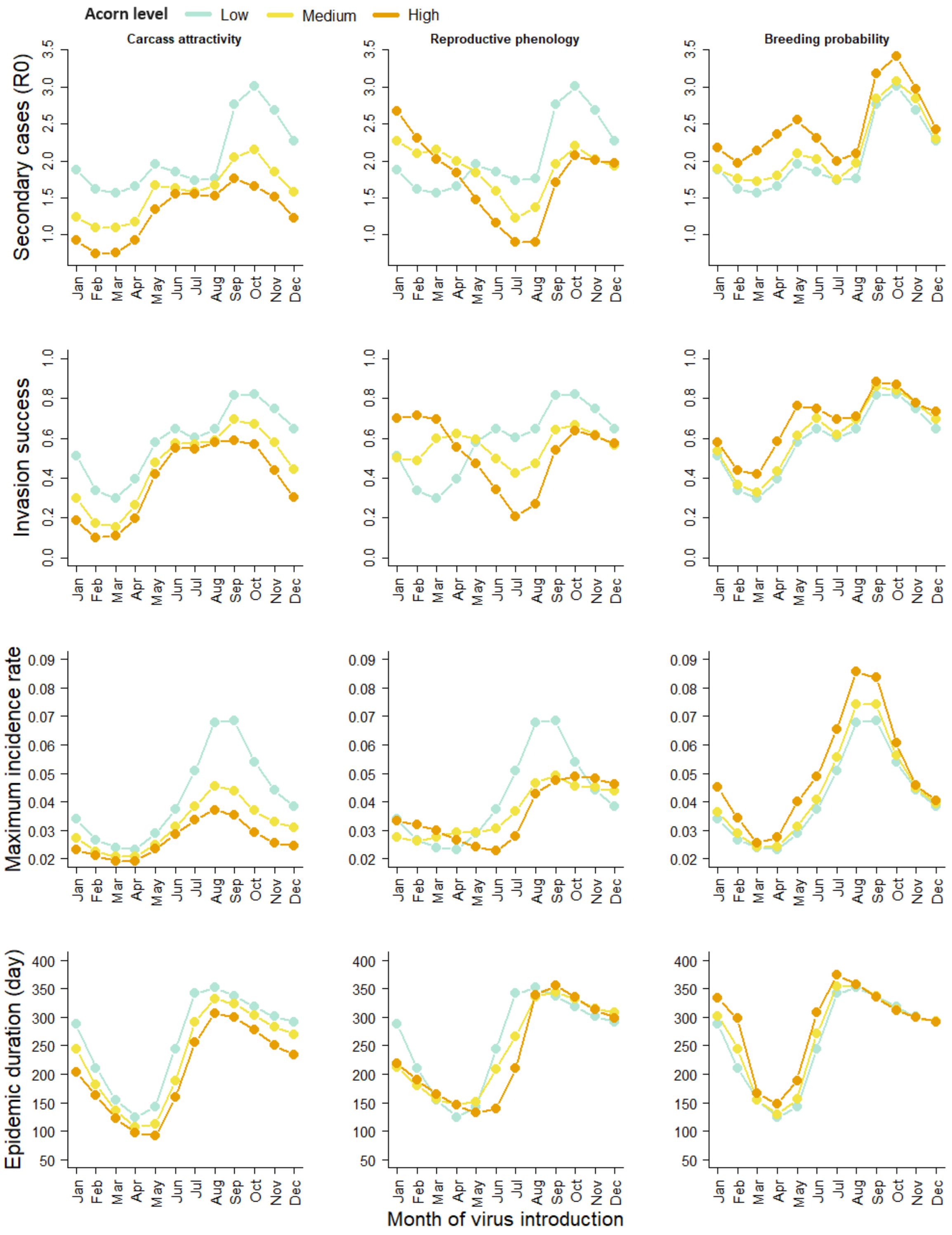

**Figure S2**: Projection of the epidemic dynamics in future climatic conditions including only direct temperature effects (i.e. influence on virus persistence in the environment). Results include (A) the basic reproduction number ($R_0$), that is the number of secondary infections generated by a single infected individual, (B) the invasion success probability defined as the probability that there were at least five secondary cases, (C) the maximum daily incidence rate and (D) the epidemic duration conditional on successful virus invasion. Results are presented as a function of time of virus introduction in the population from the 1st January to 1st December.

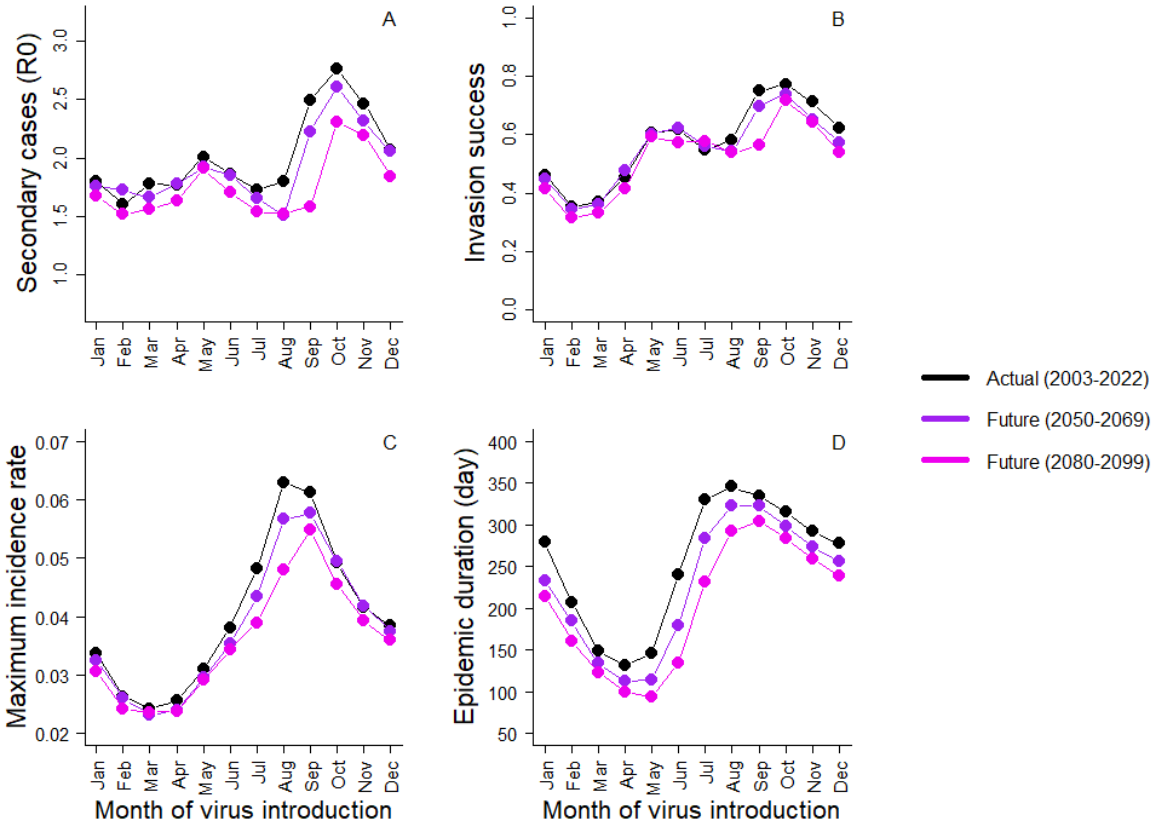

**Figure S3**: Projection of the epidemic dynamics in future climatic conditions including only indirect temperature effects (i.e. through acorn abundance). Results include (A) the basic reproduction number ($R_0$), that is the number of secondary infections generated by a single infected individual, (B) the invasion success probability defined as the probability that there were at least five secondary cases, (C) the maximum daily incidence rate and (D) the epidemic duration conditional on successful virus invasion. Results are presented as a function of time of virus introduction in the population from the 1st January to 1st December.

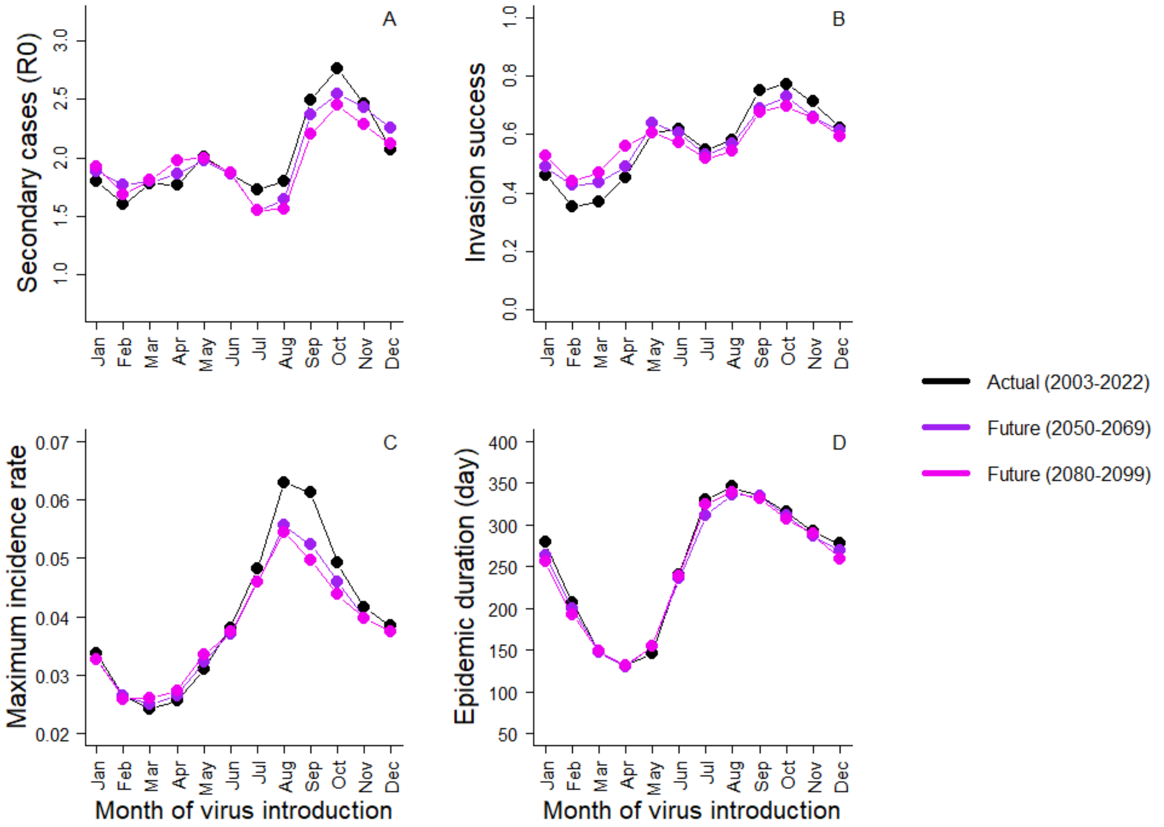

**Appendix S1**:

Purpose

This model aimed to understand the interplay between a virus (ASF) and a host population (wild boar) under variable temperature conditions. More specifically, we explored how direct and indirect temperature effects influence pathogen and host interactions, and subsequent epidemic dynamics. This model was not spatially explicit, and figured out epidemic dynamics within a closed wild boar population, corresponding to the case of a localized wild boar population, or to a population that has been isolated, as has been done in Europe following disease detection (8).

Entities, state variables, and scales

Hosts were modeled individually and characterized by sex, age and health status. The infection dynamics was modelled considering an SEIR epidemiological process (susceptible, exposed, infected and recovered). Host carcasses were characterized by weight and health status (infected or not infected). One-time step corresponded to one day and the whole simulation period covered four years (i.e., 1,460 time steps).

Process overview and scheduling

Each day, individual boar may successively survive or die, breed and change health status.

*Survival* - Survival probability varied with age, sex, health status and the period of the year. Specifically, survival probabilities were lower during the hunting season than during the non-hunting season. We also assumed that piglets less than 1-month old died if they lost their mother.

*Health status* - Health status may change from susceptible to exposed, from exposed to infected, and from infected to recovered. Transition between susceptible to exposed occurred as a function of three components: (i) the number of contacts an individual has with its relative or carcasses; (ii) the proportion of those contacts with infected individuals or carcasses; and (iii) the proportion of infectious contacts that actually resulted in infection. Exposed individuals became infectious after an incubation period of four days, and remain infectious for an average of 5 days (7). At the end of the infectious period, all infected individuals had the same mortality risk regardless of their age and sex. If the individual survived the disease course, it became immune and cannot be infected again, nor can it be infectious. No maternal immunity transfer from immunized sows to newborn piglets was assumed.

*Carcass degradation* - Infected carcasses were potential source of infection as long as they remained present in the environment. Carcass degradation was modelled as a function of their size and the air temperature.

*Reproduction* - Reproduction was decomposed in three successive steps: the reproduction probability, the litter size and the sex-ratio. The reproductive probability was variable according to the period of the year (daily resolution), and both reproduction probability and litter size vary with age (reproductive potential increase with age). Females may reproduce from subadult age and once a year at maximum, regardless of their age. Reproductive probability (both values and timing) was influenced by acorn abundance. Finally, reproduction was constraint by male presence: female reproduction was not possible if all sexually mature males (>2-year-old) disappeared. Sex-ratio was assumed balanced (6).

<u>Design concept</u>

Interaction: Each day, individuals may interact with others live individuals or with carcasses, representing opportunities for new infections. The number of contacts with carcasses was invariant across individuals but varied with acorn abundance (see sub-model below). The number of contacts among live individuals was age and sex-specific - females and young's have more contacts with their congeners. For adult males, contact rate with congeners varied seasonally being stronger during the mating season (9). Because contact rate increases typically nonlinearly with population density (10), we assumed that the frequency of contacts among live individuals increased with population size (see submodels). Beyond these factors, we assumed homogeneous mixing of individuals within this population.

Stochasticity: The realizations of wild boar daily vital rates (survival, reproduction, litter sex-ratio) were stochastic as well as the infection risk and the infection period.

Emergence: The first emerging result from our IBM was the wild boar population dynamic estimated at a daily time step, which resulted from the day-to-day survival and reproduction of the individuals. Our model provided estimations of the seasonal variation in population size and population structure (age and sex) along the year at a daily time step. The second emerging result was the epidemic dynamic (e.g. the daily number of new infected individuals, epidemic duration) that resulted from the interplay between the wild boar population dynamic and the pathogen dynamic.

Observation: The observations were done from an omniscience perspective. Each day, we observed the host population size and composition (age, sex, health state), and the number of carcasses, including their sanitary condition. We tracked the number of secondary infections caused by the first infected individual ($R_0$), and the daily number of new cases, distinguishing those caused by the two types of contacts (live vs. carcass).

<u>Initialization</u>

Each simulation run was started assuming a population of 100 wild boars. Initial age and sex distributions were attributed according to the stable age and sex distribution. All individuals were immunologically naïve (susceptible). Host population was projected one year before the introduction of the virus in order to insure independence from the initial conditions. Virus was introduced on the first of each month in the population, adding one infected sub-adult male (age = 1.5 years), mimicking the immigration of an infected individual within the population.

Input data

Empirical data were also used to estimate the average daily air temperature variation across and among the years, which affect the speed of carcass degradation (see carcass degradation submodel). We used daily air temperature collected from 2003 to 2022 on six Météo France weather stations surrounding our study area (<50km). Finally, stomach contents of shot individuals were used to estimate time series of annual acorn abundance (three categories: low, medium and high acorn abundance) from 2003 to 2022.

Submodels

Parameters used in the submodels are summarized in Table S1.

**Wild boar survival:** Survival was modelled using a Bernoulli distribution. Age-specific survival probabilities were derived from size-specific survival estimates using published sex-specific growth trajectories (11). Daily survival probabilities were obtained assuming constant survival rate over the hunting season (15 Oct. to 15 Feb., 241 days) and over the non-hunting season (16 Feb. to 14 Oct., 124 days). Note that survival probability was influenced by the disease (see virus component).

**Wild boar reproduction:** Reproduction was decomposed in three successive steps: the breeding probability, which was modelled using a Bernoulli distribution, the liter size, which was modelled using a multinomial distribution, and the piglet sex-ratio, which was balanced in average but randomly determined with a Binomial distribution. Daily breeding probabilities were derived from the annual breeding probabilities and the birth distribution. Both breeding phenology and breeding probabilities were influenced by acorn abundance, distinguishing three set of parameters corresponding to the three levels of acorn production (low, medium and high) (see Fig. 1 and Table S1). We used the observed litter size distribution to determine the probability of occurrence of each litter size. As for survival, size-specific reproductive parameters were converted to age-specific parameters using the mean female growth trajectory (11).

**Infection risk**: Infection risk was modelled using a Binomial distribution where the number of trials correspond to the number of infectious contacts and the success probability corresponded to the risk to be infected per infectious contact. The number of infectious contacts was equal to the daily number of contacts multiplied with the proportion of infectious contacts (i.e., proportion of sick individuals or infected carcasses). The infection risk per infectious contact vary according to the type of contact. In the two following paragraphs, we describe how the number of contacts with other live individuals and carcasses was computed.

**Contact with live individuals**: The daily number of contacts per individual was modelled as function of population size, age, sex and for reproducing males, day. We first used a generalized logistic model to simulate density dependent contact numbers:

$$DC_0 = A + \frac{K - A}{(1 + e^{-rN})^{1/v}}$$

where *N* is the total population size, *A* and *K* are the lower and higher asymptotes respectively, *r* is a growth rate and *v* affects near which asymptote maximum growth occurs. Second, we corrected this daily number of contacts using age- and sex-specific factors, and a function allowing to account for the seasonality of contacts for sexually mature males:

$$DC_{i,t} = \frac{DC_0}{\gamma_{age_i} + \gamma_{sex_i}} \times f_{t,sex_i}$$

where *i* and *t* are indices for individuals and days respectively, $\gamma_{age}$ have three levels corresponding to juvenile individuals (< 6 months), sub-adults (6 months to 2 years) and adults (> 2 years), and $\gamma_{sex}$ has two levels, one per sex. We assumed that sex effect was null for the youngest individuals (< 6 months), because both sexes live within social groups at this early age (9). $f_{t,sex}$ is a function which modulates the number of contacts according to the day *t* of the year, and applies for old males only (> 2 years). This function allows modelling males behavior change during the rut from a solitary to a harem mode of life (9).

**Contact with carcass**: We assumed that the number of contacts with carcasses per individual was frequency dependent (i.e., the number of contacts is proportional to the total number of carcasses in the environment). The daily number of contacts with carcasses per individual was modelled using the following linear model:

$$IC = \delta \times NC$$

Where *IC* is the daily number of contacts with carcasses per individual, *NC* is the total number of wild boar carcasses and $\delta$ the rate of increase of contact per unit of carcasses available in the environment. We assumed that $\delta$ decreases with acorn abundance (Figure 1B), hence reflecting the preference of wild boars for acorn and their increasing foraging activity when this resource is rare, which is expected to increases the probability to cross and contact a carcass (12–14). Quantitative measurements of these changes in contact rates are inexistent in the literature. Thus, to be conservative, we parametrized our model using a modest decrease of carcass attractivity with acorn abundance. We assumed that contacts with carcasses were four times less frequent when acorn abundance shift from low to high (Table S1).

**Disease course**: In case of exposure, after a fixed incubation period of four days, the infection period was determined stochastically using a one truncated Poisson distribution with a mean of five. Thus, the modal infection period was 5 days and 98% of infection periods range between 1 and 9 days.

**Carcass degradation**: Carcass degradation was modeled as a cumulative process. We defined a degradation coefficient that varied from 0 (no degradation corresponding to the day of death), to 1 (full degradation). Each day, the degradation coefficient of a carcass was updated adding the degradation occurring since the previous day. The daily degradation (DD) of the carcass *i* on day *t* was computed as a function of the daily mean air temperature $T_t$ (°C) and carcass mass at death $M_i$ (Kg) using the following equation:

$$DD_{i,t} = \frac{1}{(-4.076 \times T_t + 92.75) \times (-0.0143 \times M_i + 1.572)}$$

We parametrized this model to mimic boar carcass degradation speed observed empirically (13, 15). Under this model, the full degradation of a 40 kg carcass takes around two to three weeks in summer and three months in winter, and these durations increase with carcass size being roughly increase by 33% for carcass of 80 kg.
